# Immunomodulatory Porous Regenerative Scaffolds for *in situ* Vascular Engineering

**DOI:** 10.1101/2024.01.29.577757

**Authors:** Le Zhen, Elina Quiroga, Sharon A. Creason, Ningjing Chen, Tanmay R. Sapre, Jessica M. Snyder, Sarah L. Lindhartsen, Brendy S. Fountaine, Michael C. Barbour, Syed Faisal, Alberto Aliseda, Brian W. Johnson, Jonathan Himmelfarb, Buddy D. Ratner

## Abstract

The 70-year quest for synthetic vascular graft (sVG) endothelialization has not led to completely healed endothelium in clinically used sVGs. In humans, healing is limited to the vicinity of anastomotic regions (pannus ingrowth) and does not reach the middle regions of sVGs. Here, we conducted proof-of-concept implantation of *immunomodulatory porous regenerative scaffolds for in situ vascular engineering* (IMPRESSIVE) as interposition grafts in sheep carotid arteries. These scaffolds are based on a new polyurethane (PU) material featuring a 40 µm precision porous structure optimized for angiogenesis. The modulus of the PU was adjusted to match that of natural arteries. The implantation study revealed rapid healing in IMPRESSIVE sVGs. In side-by-side comparison with standard polytetrafluoroethylene (PTFE) grafts, the luminal surfaces of PU grafts were almost completely covered with nucleated cells, while healing in PTFE grafts was limited to several millimeters within anastomotic regions. Endothelialization was observed in the middle regions of PU grafts and overall endothelialization increased significantly compared to PTFE grafts. Densities of mononuclear cells, foreign body giant cells (FBGCs), and endothelial cells within graft walls of PU grafts were also significantly higher than those in PTFE grafts, suggesting transmural cellular infiltration may play a key role in overall improved healing. High percentages of macrophages in pores of PU grafts show Type 1 (CCR7+) and Type 2 (mannose receptor, MR+) characteristics. We also discovered that FBGCs exist in a diverse spectrum of phenotypes. Dually activated FBGCs (CCR7+MR+, G1/2) dominate the population of FBGCs associated with pro-healing PU grafts. These observations suggest a complex, balanced pro-healing response from macrophages and FBGCs. The IMPRESSIVE approach may enable complete endothelialization in pro-healing sVGs and have wide applications in implantable devices and tissue engineering.

## Introduction

Since Dr. Arthur B. Voorhees and colleagues reported the first functioning synthetic vascular graft (sVG) in 1952^1^, the healing of sVGs, especially the recapitulation of the endothelium (the luminal lining of native blood vessels), has been the “holy grail” for vascular graft development. However, the 70-year+ quest has not led to completely healed sVGs in clinical settings. The consistent observation in humans is that tissue heals from the ends (anastomosis) of sVGs where they are sutured to native blood vessels (BVs) and stops after several millimeters of outgrowth. This is called trans-anastomotic ingrowth, or pannus ingrowth, which can only heal near the anastomosis. The middle region of sVGs is always devoid of healed tissue.^2^

*In vitro* tissue engineering offers an exciting, interdisciplinary approach to produce blood vessels from living cells.^3,4^ However, complex *in vitro* procedures, long production time, high cost, problems from cell sourcing and expansion, and concerns with host compatibility slow the translation of such promising technologies into the clinic. To implant such engineered blood vessels into patients, they must be decellularized, leaving behind biological scaffolds for the host tissue to heal into.^5^ Direct engineering of synthetic, regenerative scaffolds that recruit host cells *in situ* to reconstruct new tissue (*in situ* tissue engineering) provides a path that circumvents the complexity of *in vitro* tissue engineering bioreactors and can lead to rapid clinical translation.

Porosity (here defined as conditions and qualities of being porous) has been found since the time of Arthur Voorhees to be important for sVG performance. Even in the early days of sVG research, it was observed that, with sufficient porosity (characterized by water permeability), sVGs perform relatively well, seemingly regardless of the material used to construct the sVGs.^6^ In expanded poly(terafluoroethylene) (ePTFE) grafts, the most widely used sVGs, it was observed that, when porosity (characterized by internodal distance) was increased from the standard 30 µm to 60 µm and above, tissues (blood vessels, in particular) are able to heal through the graft wall (transmural ingrowth) and reach the lumen of the middle region of sVGs.^7^ However, in order for sufficient durability for clinical implantation, ePTFE grafts needed to be externally reinforced, which likely impeded the transmural ingrowth in a clinical trial.^8^ Although the importance of porosity is well-recognized, pore structures in sVGs are seldom precision controlled, impeding the elucidation of the optimal pore size for healing.

Our laboratory developed precision porous materials where all pores are spherical and interconnected, and we discovered (through implantation in small animal models) that the optimal pore size for endothelial cells recruitment and tissue reconstruction is around 35-40 µm.^9–11^ The precision porous materials likely achieve the pro-angiogenic effect by attracting macrophages and modulating them towards a complex, pro-healing phenotype.^10^ We optimized the mechanical properties of this pore structure with a new biostable, elastic, precision porous scaffold with mechanical properties matching those of native blood vessels, and verified the pro-angiogenic effect through implantation in mice.^12^ Whether such a scaffold can enable *in situ* vascular engineering and lead to healed vascular grafts in a clinically relevant large animal model was assessed here.

An appropriate animal model is crucial for the assessment of sVG healing. Complete healing is frequently reported in many animal model studies, yet never realized in adult humans. One problem may be that the animal models used, such as mice,^13^ rats,^14^ dogs,^15^ pigs,^16^ and even baboons^5,7^, most often young animals, are intrinsically better at healing compared to adult humans. Adult sheep have emerged as a large animal model that recapitulates the limited healing capacity in humans.^16,17^ Such a model can generate clinically relevant results for sVG healing.

In the presence of foreign material, such as an sVG, some macrophages fuse to form multinucleated giant cells (MNGCs). These MNGCs closely associated with foreign materials are also known as foreign body giant cells (FBGCs). FBGCs are widely considered the “villain” in the body’s response to foreign body (FBR), because they are implicated in implant degradation and failure.^18^ FBGCs are also thought to be terminally differentiated,^19^ thus may not have the phenotypical plasticity of macrophages. However, our previous study shows large numbers of FBGCs associated with the pro-healing response of the precision porous material.^12^ And a diverse spectrum of phenotypes between classically activated (Type 1, “pro-inflammatory”, M1) and alternatively activated (Type 2, “pro-healing”, M2) are exhibited in macrophages, the cells FBGCs originate from.^20^ Based on these observations, a re-examination of the role of FBGCs in the FBR and their phenotypic diversity is necessary.

In this study, we conducted a proof-of-concept implantation of *immunomodulatory porous regenerative scaffolds for in situ vascular engineering* (IMPRESSIVE) in sheep. These scaffolds are made of a new polyurethane (PU) elastomer optimized for sVG applications.^12^ We hypothesize that the IMPRESSIVE PU grafts improve healing by recruiting macrophages and FBGCs, and modulating them towards complex and pro-healing phenotypes.

Compared to small, elongated, irregular pores in standard PTFE graft, the IMPRESSIVE PU graft features a precision-controlled porous structure optimized for angiogenesis.^9–12,21^ The PU graft also has a higher compliance more closely matching that of natural artery and leads to less disturbance of blood flow. PU grafts were implanted side by side with standard PTFE grafts in bilateral interposition grafting in carotid arteries. Using PTFE grafts as the control, we observed that healing is limited to several millimeters within the anastomosis in PTFE grafts, confirming the relevance of the model to human. In contrast, PU grafts are almost completely cellularized. In addition, both the density of endothelial cells in pores and the coverage of endothelial cells on the lumen (endothelialization) are significantly increased in PU grafts. Importantly, endothelialization at the middle region of the lumen of PU grafts was observed. While the interstices of PTFE grafts are almost completely filled with red blood cells (blood clot), the pores throughout the entire graft wall of PU grafts are infiltrated with nucleated cells and blood vessels. Among the cells are large numbers of macrophages, high percentages of which exhibit M1 and M2 characteristics. PU grafts also attracted higher density of FBGCs compared to PTFE grafts. We also discovered diverse phenotypes of FBGCs, including unactivated (G0), classically activated (G1), alternatively activated (G2), and dually activated (G1/2). The majority of FBGCs associated with PU grafts are G1/2, indicating G1/2 to be a complex, pro-healing phenotype. The IMPRESSIVE approach provides an opportunity to rapidly heal vascular grafts. This approach is readily translatable to the clinic because *in vitro* pre-treatment is not needed. In addition, the observation of a diverse spectrum of FBGC phenotypes may fundamentally change the understanding of their potential functions in the FBR and biocompatibility.

## Results

### The porous structure of PU grafts is precise and permissible for cell infiltration

To compare the microporous structure of PTFE and PU grafts, scanning electronic microscopy (SEM) was used to examine the cross sections (**Fig. 1**, first five rows), exterior surfaces (**Fig. 1**, the sixth row), and luminal surfaces (**Fig. 1**, the last row) of both grafts. Low and medium magnification SEM images of the whole graft wall cross sections reveal that the PU graft wall has a uniform, precision controlled, interconnected porous structure, while the PTFE graft wall is bi-layered consisting of a primary wall structure and an exterior reinforcement structure (**Fig. 1**, 1^st^ & 2^nd^ row). The high magnification image of luminal side of the PTFE graft cross section shows that the primary wall structure consists of layers of long strips of non-porous PTFE (called “nodes”, extending on the circumferential direction) interconnected with paralleled fibrils running on the longitudinal direction of the graft (**Fig. 1**, 3^rd^ row, left).

**Figure 1.**
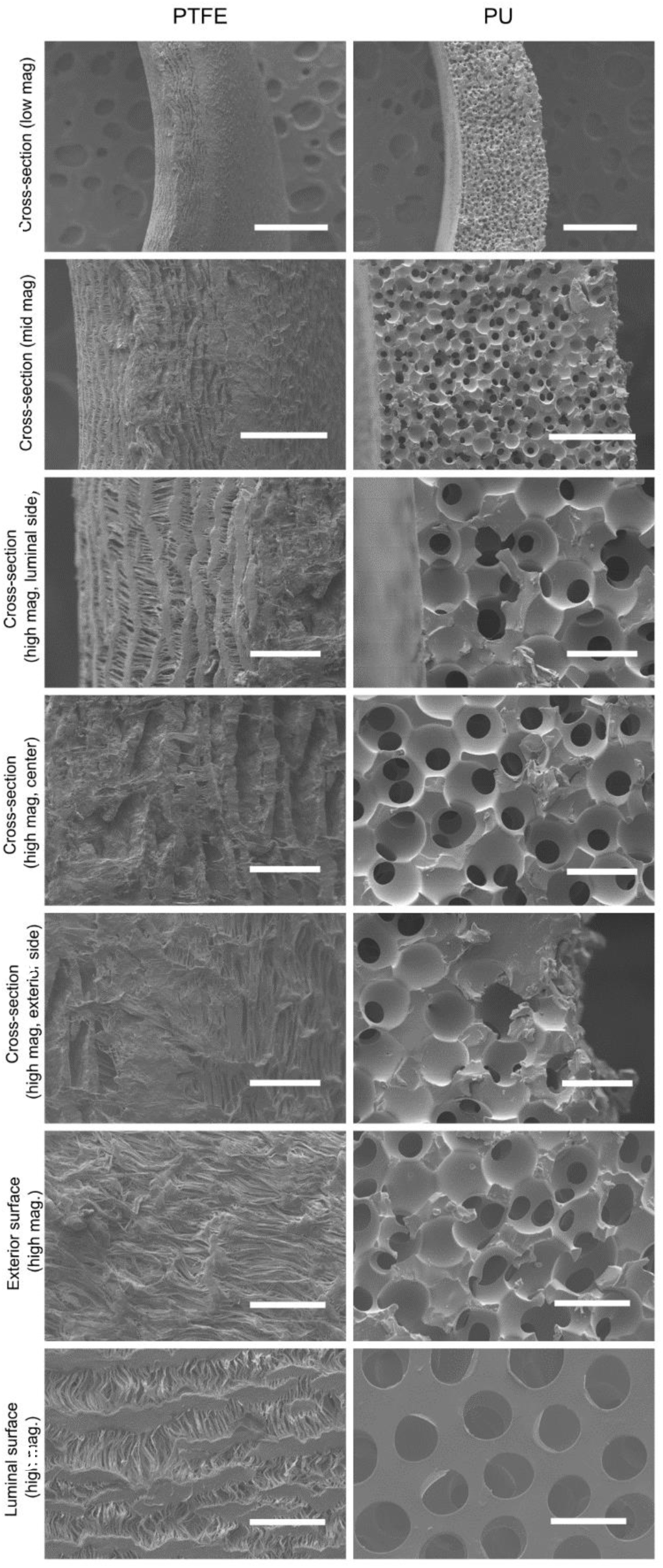
SEM images of microporous structures of PTFE (first column) and IMPRESSIVE PU grafts (second column). Low magnification (100×) scale bar: 500 µm; mid-magnification (300×) scale bar: 200 µm; high magnification (1000×) scale bar: 50 µm.

Many layers of such structures irregularly stack together to form the primary graft wall (**Fig.** 1, 4^th^ row, left). Such a structure was generated through a heated extension process along the longitudinal direction, thus the name expanded PTFE. The exterior surface of the PTFE graft is reinforced with a thin expanded PTFE film with much smaller nodes (aligned along the longitudinal direction), and much more densely aligned fibrils (along the circumferential direction), resulting in much less porous structure compared to the primary wall (**Fig. 1**, 5^th^ row, left). The porous structures of the exterior reinforcement and the luminal surface are better visualized in *en face* images (**Fig. 1**, last two rows, left). Even under 1000× magnification, few open pores can be seen on the exterior surface. The porous structure on the luminal surface is more open, characterized by an average internodal distance (distance between the mid-lines of two adjacent nodes connected by fibrils) of 30 µm.^7^ However, the sizes of pores are also limited by the distances between adjacent fibrils, which are mostly below 5 µm (about half of the size of the nucleus in a typical mammalian cell). On the other hand, the porous structure of the PU graft is consistent throughout the graft wall (**Fig. 1**, 3^rd^ to 5^th^ row, right). Such a porous structure is characterized by a spherical pore diameter of 40.8 ± 0.4 µm, and an interconnection diameter of 18.0 ± 0.7 µm. The porous structure on the exterior surface is identical to that in the cross section (**Fig. 1**, 6^th^ row, right). The luminal surface is relatively smooth and punctuated by an array of holes connecting to the porous structure underneath (**Fig. 1**, the last row, right). The diameter of the holes is 27.7 ±1.2 µm. The porous structure of the PU graft is optimized for cell infiltration and angiogenesis.^10–12^ To prevent hemorrhage immediately after implantation, the pores of PU grafts were sealed with a slightly crosslinked gelatin sealant that completely degrades within 3 weeks *in vivo* (**Fig. S1**). Overall, the porous structure of PU grafts is precision-controlled and permissible for cell infiltration, while the porous structure of PTFE grafts varies substantially and would likely prohibit cell infiltration due to size exclusion.

### Physical measurements reveal multiple favorable properties of PU grafts

The compliance of grafts was measured using a custom-made dynamic flow system (**Fig. S2**) in accordance with ISO 7198.^22^ A representative, real-time diameter (top), pressure (bottom) profile of a PU graft (**Fig. 2a**, left) shows synchronized, pulsatile pattern of diameter and pressure versus time. Compliances calculated from such profiles show that PU grafts (compliance: 2.40±0.32 %/100 mm Hg) are about 8 times more compliant than PTFE grafts (compliance: 0.27±0.03 %/100 mm Hg) over the pressure range of 50-150 mm Hg (a pressure range sufficient to generate measurable diameter changes for both grafts). The compliance of PU grafts was further measured in three pressure ranges (**Fig. 2a**, right): 50-90 mm Hg (compliance: 2.82±0.18 %/100 mm Hg), 80-120 mm Hg (compliance: 2.25±0.13 %/100 mm Hg), 110-150 mm Hg (compliance: 1.86±0.27 %/100 mm Hg). The trend of decreasing compliance with the increase of pressure is consistent with the behavior in native blood vessels.^23^ In addition, both PU and PTFE grafts are able to withstand an internal pressure of at least 300 mmHg. Observations of grafts immediately before and after implantation also reveal desirable properties of PU grafts (**Fig. 2b**). The PU graft can be securely sutured onto the native blood vessel with the reinforcement mesh within the graft wall clearly visible, and without hemorrhage (**Fig. 2b**, left). No suture failure occurred in any of the six PU grafts after the surgery. A tight binding between the PU graft and the surrounding tissue was established after the implantation, demonstrated by the surgeon pulling on tissue partially dissected away from the graft (**Fig. 2b**, 2^nd^ to the left). The most dramatic difference between the PU graft and the PTFE graft was observed on the luminal surface, where the PU graft (top) was covered with a thin layer of tissue with very few blood clots (red) attached, while the PTFE graft was almost completely covered with red, blood clots with little healed tissue (**Fig. 2b**, 2^nd^ to the right). This strongly suggests the PU grafts’ improved healing and its potential beneficial effect on blood compatibility. To further examine such healing histologically, grafts together with the surrounding tissue were trimmed (**Fig. 2b**, right), embedded in paraffin, and sectioned (along the red lines).

**Figure 2.**
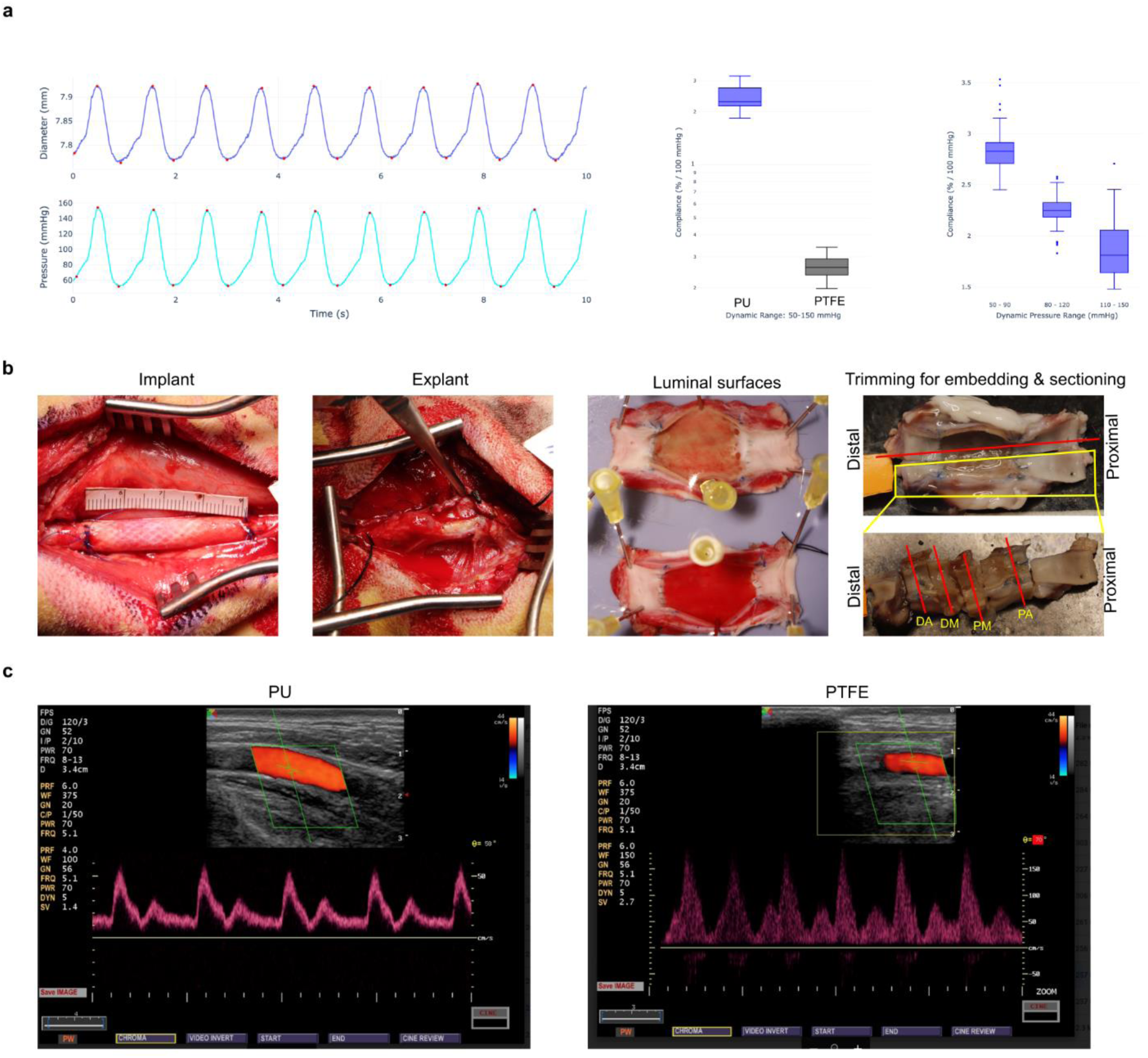
Physical observations on vascular grafts. **a)** Compliance measurements: (left) representative data showing the pulsatile behavior in the dynamic flow system (dynamic pressure range: 50-150 mmHg); (middle) compliance (dynamic pressure range: 50-150 mmHg) of PU and PTFE grafts; (right) compliance of a representative PU graft over different dynamic pressure range (50-90 mmHg, 80-120 mmHg, 110-150 mmHg). **b)** Gross observations of graft-tissue interactions: (left) a PU graft immediately after sutured into the carotid artery (a 4-cm-long ruler was placed next to the graft for scale); (second to the left) a PU graft integrated with surrounding tissue at explant (the surgeon was pulling on the surrounding tissue to demonstrate the tight binding); (second to the right) a side-by-side comparison of luminal surfaces of PU (top) and PTFE (bottom) grafts implanted bilaterally in the same sheep; (right) a representative image demonstrating the trimming of a graft in preparation for histological sectioning. (red lines indicate the direction of sectioning; PA: proximal anastomosis; PM: proximal mid-graft; DM: distal mid-graft; DA: distal anastomosis.) **c)** Representative ultrasound images of PU (left) and PTFE (right) grafts taken before explant.

Doppler ultrasound imaging was conducted before each retrieval surgery to examine the patency of grafts. Although all grafts remains patent over the implantation period, ultrasound spectral broadening was observed in PTFE grafts, not in PU grafts (**Figure 2c**), indicating more turbulent flow in PTFE grafts.^24,25^ This may be attributed to flow disturbances caused by the low compliance of PTFE or the relatively uneven lumen of the PTFE grafts (**Fig. 1**). All observations and measurements suggest that the PU grafts are more compliant, have better biointegration, reduce adherent thrombus and produce more physiological flow patterns.

### Basic histology shows improved biointegration and reduction of calcification for PU grafts

To investigate biointegration, whole graft histological sections were stained with H&E and Masson’s trichrome. H&E stained sections show that the lumen of the PU graft is mostly covered with tissue, while tissue coverage in the PTFE graft is limited to the vicinity of anastomoses (**Fig. 3a**). Trans-anastomotic ingrowth, or pannus ingrowth, can be distinguished by a continuous decrease in tissue thickness towards the center of the grafts (**Fig. 3b**, arrows point at the end of pannus ingrowth). On the other hand, mid-graft tissue (most likely from transmural ingrowth) is relatively thin and uniform in thickness (**Fig. 3b**). Quantitative analysis of the length of pannus ingrowth reveals almost identical behavior in PU and PTFE grafts--the ingrowth from the proximal end (∼ 2 mm) is slightly more advanced than that from the distal end (∼ 1.5 mm), but far from reaching the mid-graft region (**Fig. 3c**, top left). In fact, the total luminal tissue coverage of PTFE grafts (consisting entirely of pannus ingrowth) is only 14 ± 3 %, while this value rises to 95 ± 4 % in PU grafts (**Fig. 3c**, top right, p=7.8×10^-8^), suggesting that transmural ingrowth likely contributes to most (∼80 %) of the luminal tissue coverage in the PU graft.

**Figure 3.**
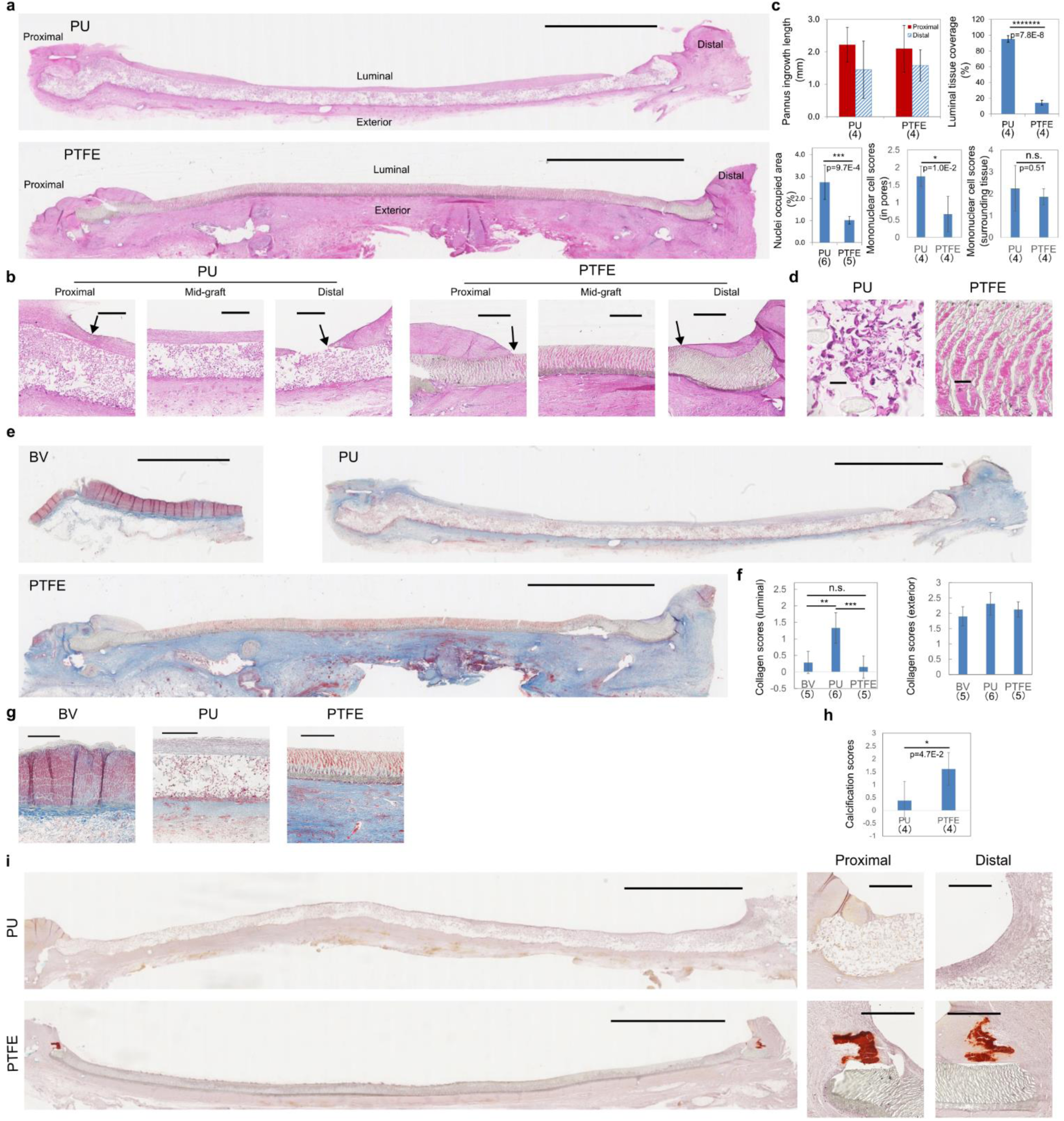
Basic histological stains: H&E (**a-d**), Masson’s trichrome (**e-g**), and Alizarin Red S (**h, i**). **a)** Low magnification micrographs of the entire H&E stained, longitudinal section of PU (top) and PTFE (bottom) grafts; (scale bars: 5 mm; the orientations of following longitudinal sections are consistent as labeled here.) **b)** mid-magnification micrographs of selected regions (proximal, mid-graft, distal) of PU (left 3 images) and PTFE (right 3 images) grafts; (scale bars: 500 µm; arrows point at the end of pannus ingrowth.) **c)** quantitative and semi-quantitative analysis of H&E stained grafts; **d)** high magnification micrographs of H&E stained tissue in porous structures of PU (left) and PTFE (right) grafts (mid-graft region); (scale bars: 50 µm) **e)** Low magnification micrographs of Masson’s trichrome stained, longitudinal section of a native blood vessel (BV, top left), PU (top right) and PTFE (bottom) grafts; (scale bars: 5 mm) **f)** semi-quantitative scores for collagen capsule on the luminal side (left) and the exterior side (right); **g)** mid-magnification micrographs of BV, PU and PTFE grafts (mid-graft region); (scale bars: 500 µm) **h)** semi-quantitative scores of Alizarin Red S stained PU and PTFE grafts; **i)** Representative Alizarin Red S stained PU (top) and PTFE (bottom) grafts, where low magnification (scale bars: 5 mm) micrographs of full length longitudinal sections are on the left, and further magnified micrographs (scale bars: 500 µm) of the anastomosis region are shown on the right. Numbers of biological replicates are shown in parentheses under group labels in bar graphs.

The extent of transmural ingrowth was further quantified by the density of nuclei within graft walls (**Fig. 3c**, bottom left). The nuclei density in the wall of PU grafts is significantly higher than that of PTFE grafts (p=9.7×10^-4^). This is confirmed by high resolution images of the graft walls (**Fig. 3d**). The pores of the PU graft are densely occupied by nucleated cells (**Fig. 3d**, left), while the interstices of the PTFE are almost entirely occupied by red blood cells (blood clot, **Fig. 3d**, right), but not blood vessels. Rigorous rinsing with heparinized saline after graft harvesting ensured the removal of liquid blood, which strongly suggests that the large number of red blood cells left in PTFE grafts are a part of solid clot. There is intense transmural healing in PU graft walls, and almost no healing in PTFE graft walls. This is consistent with the observation that the pore size of PU grafts is permissive of nucleated cell infiltration, while that of PTFE grafts is likely prohibitive. The density of mononuclear cells (mainly macrophages) is significantly higher in PU graft walls (**Fig. 3c**, bottom middle, p=0.01), but remains comparable in the surrounding tissue of both grafts (**Fig. 3c**, bottom left, p=0.51). This suggests a role of mononuclear cells infiltrating the graft wall in promoting healing.

To examine collagen deposition, both grafts along with native blood vessels (BV, carotid artery) were stained with Masson’s trichrome (**Fig. 3e**-**g**). Images of a BV (**Fig. 3e**&**g**) show that the BV wall consists of red, densely packed smooth muscle cells, and is surrounded by a dense layer of collagen (dark blue) that histologically resembles the classic FBC observed in the subcutaneous region of mice.^12^ This dense adventitial collagen layer represents the “normal” collagen deposition around BVs. In addition, diffuse collagen (light blue) is occasionally observed on the luminal surface (**Fig. 3g**, left). PU and PTFE grafts are also surrounded by relatively dense collagen layers (**Fig. 3e**&**g**). The lumen of the PU graft is mostly covered by a thin layer of diffuse collagen, while the luminal collagen coverage of the PTFE graft is limited to the vicinity of anastomoses (**Fig. 3e**&**g**). This observation mirrors that of the luminal tissue coverage. Collagen depositions on the luminal side and the exterior side were separately scored by a pathologist blinded to the hypothesis of the study (**Fig. 3f**). This scoring system depicts collagen deposition in terms coverage, density, and thickness.^12^ PU grafts receive the highest score (1.3±0.5) on the luminal side, which is significantly higher than BVs (0.3±0.3) and PTFE grafts (0.2±0.3). This is consistent with the observation that only the lumen of PU graft is mostly covered with diffused collagen. This collagen layer may provide an ideal substrate for endothelial cells to grow on. On the exterior side, the collagen scores received by all three groups are around 2 and statistically indistinguishable (**Fig. 3f**, right, F ratio=2.25< F(critical)=3.80). This shows that the collagen layers surrounding both PU and PTFE grafts are comparable to the normal collagen layer surrounding BVs. The collagen deposition surrounding the grafts may serve as a reinforcement.

Mineralization was noted on H&E stained sections of some grafts, a sign of calcification. To assess the level of calcification in grafts, whole graft histological sections were stained with Alizarin Red S (**Fig. 3i**) and scored by a pathologist blinded to the hypothesis of the study (**Fig. 3h**). PTFE grafts receive significantly higher calcification scores than PU grafts (pairwise t-test, p=0.047, **Fig. 3h**), indicating more severe calcification in PTFE grafts.

Representative images of Alizarin Red S stained grafts (**Fig. 3i**) show no calcium deposits in the PU graft (top), and clear calcium deposits (red) at the anastomosis regions of the PTFE graft (bottom). This observation is consistent with the overall calcification scores of the grafts.

The improvements in cell infiltration, luminal tissue and collagen coverage, and the reduction in calcification demonstrate the PU graft’s integration with local tissue. Such integration should reduce undesirable outcomes such as calcification.

### PU grafts improve endothelialization

To assess endothelialization on both grafts, whole graft histological sections were stained with anti-CD 31 antibodies (blue, **Fig. 4a**&**b**), and the extent of endothelialization was quantified (**Fig. 4c**). The representative whole PU graft section (**Fig. 4a**, top) shows a mostly healed lumen (arrows pointing at different regions further magnified below), consistent with the histological observations. High magnification images (**Fig. 4a**, bottom) of different regions of the lumen, including regions close to the anastomoses and mid-graft regions, show CD 31 positive (blue, indicated by arrows) endothelial cells lining the luminal surfaces in all these regions. In addition, tissue inside the pores of the PU graft is also filled with endothelial cells (**Fig. 4a**, bottom, second to the left). The representative whole PTFE graft section (**Fig. 4b**) shows luminal tissue healing confined to the vicinity of anastomoses. High magnification images (**Fig. 4b**, bottom) show that the mid-graft regions are devoid of tissue and endothelial cells (blue, indicated by arrows). This observation confirms that the collagenous and cellularized tissue covering the luminal surface is a good substrate conducive to endothelial cell growth. The interstices of PTFE material also show no sign of endothelial cell infiltration. The extent of endothelialization was quantified by measuring the length of continuous endothelial cell coverages from the proximal end (proximal), those from the distal end (distal), and those apparently not connected to either ends (mid-graft, **Fig. 4c**, top). In all three categories, as well as total coverage (sum of the three categories), PU grafts show significantly increased endothelialization compared to PTFE grafts (**Fig. 4c**, top). PU grafts (53% covered), on average, show about 250% increase in luminal coverage of endothelium compared to PTFE grafts (15% covered, **Fig. 4c**, middle, p=0.013). The endothelial cell density inside the graft wall of PU grafts is also significantly higher than that of the PTFE grafts (**Fig. 4c**, bottom, p=0.0062). The rigorous transmural ingrowth of endothelial cells in PU grafts may provide an abundant source that contributes to the increase in luminal endothelialization.

**Figure 4.**
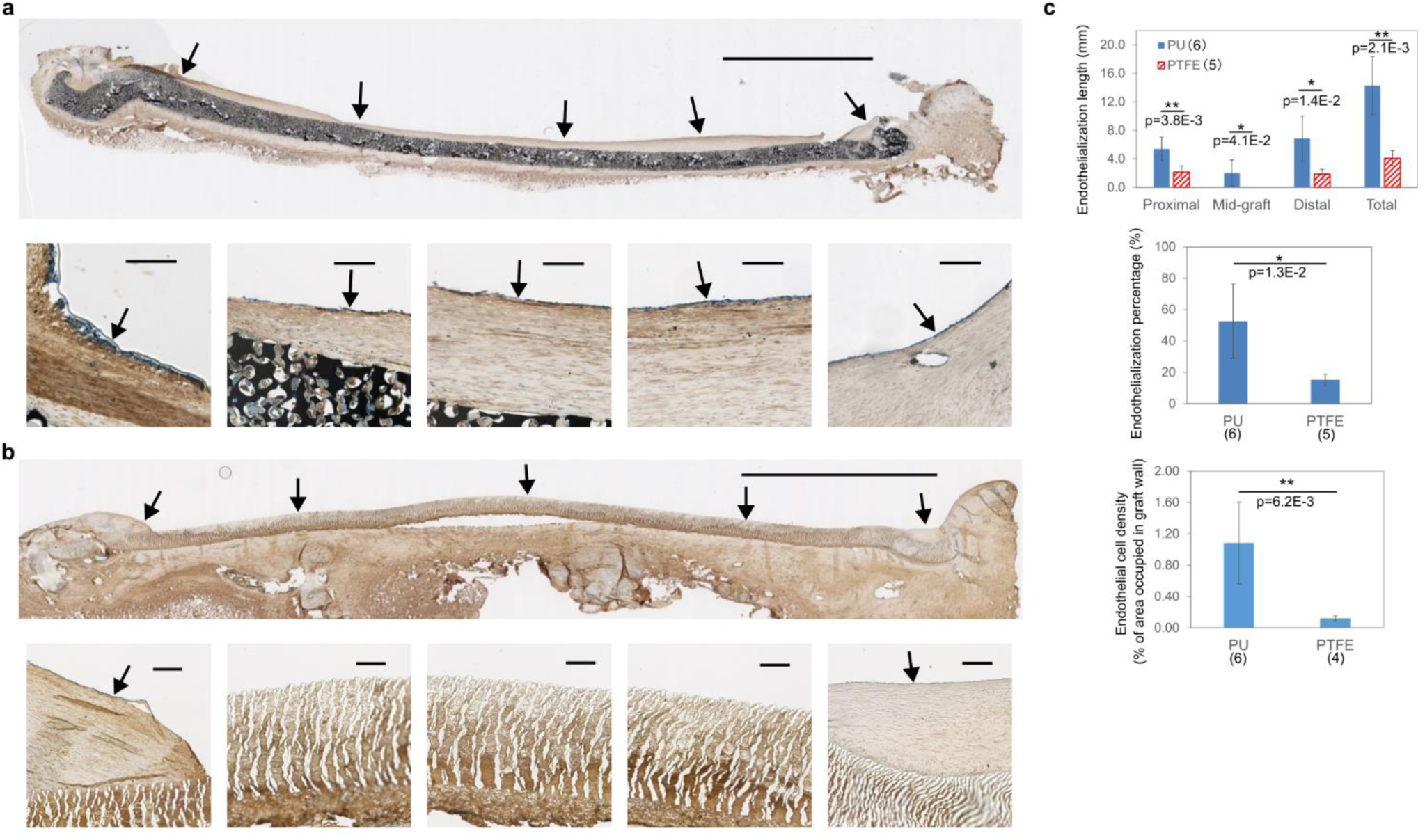
Immunohistochemistry analysis of grafts endothelialization. **a)** Representative micrographs of CD31 (in blue) stained PU graft (PU material stained in black), where the low magnification, full-length longitudinal section is shown on the top (scale bar: 5 mm, arrows point at areas shown in higher magnification below), and high magnification images of different regions of the luminal surface are shown at the bottom (arrows point at CD 31+ endothelialized lumen, scale bars: 100 µm); **b)** Representative micrographs of CD31 (in blue) stained PTFE graft (panel arrangement and labels meanings are the same as in panel **a)**); **c)** Quantitative assessment of endothelialization and endothelial cell growth: (top) Lengths of endothelialized luminal surfaces (proximal: continuous CD31+ length measured from the proximal end; mid-graft: CD31+ length apparently isolated from both ends; distal: continuous CD31+ length measured from the distal end; total: the total CD31+ length); (middle) endothelialization percentages (total endothelialized length as a percentage in total graft length) of PU and PTFE grafts; (bottom) endothelial cell density (measured by percentage of area occupied by CD31+ cells) in graft walls of PU and PTFE grafts. Numbers of biological replicates are shown in parentheses next to or under group labels in bar graphs.

### PU grafts modulate macrophages to both high M1 and M2 phenotypes

Macrophages are essential immune cells that orchestrate healing. Their phenotypic diversity is key to understanding their potential pro-healing function. To identify macrophages, circumferential histological sections were stained with macrophage specific antibody CD 64 (red, **Fig. 5a**&**c**). To identify Type 1 macrophages (M1), sections were also stained with CCR7 (blue, **Fig. 5a**) and cells positive with both CD 64 and CCR7 were identified as M1 (purple, **Fig. 5a**). Large amount of specifically stained M1 macrophages can be observed inside the pores of the PU graft (**Fig. 5a**, top). Few specifically stained cells were observed inside the PTFE grafts (**Fig. 5a**, bottom), due to the lack of nucleated cell infiltration and the abundance of nonspecific staining from blood elements occupying the interstices of the PTFE material. Clear and specific staining is observed in healed tissue surrounding both grafts (**Fig. 5a**).

**Figure 5.**
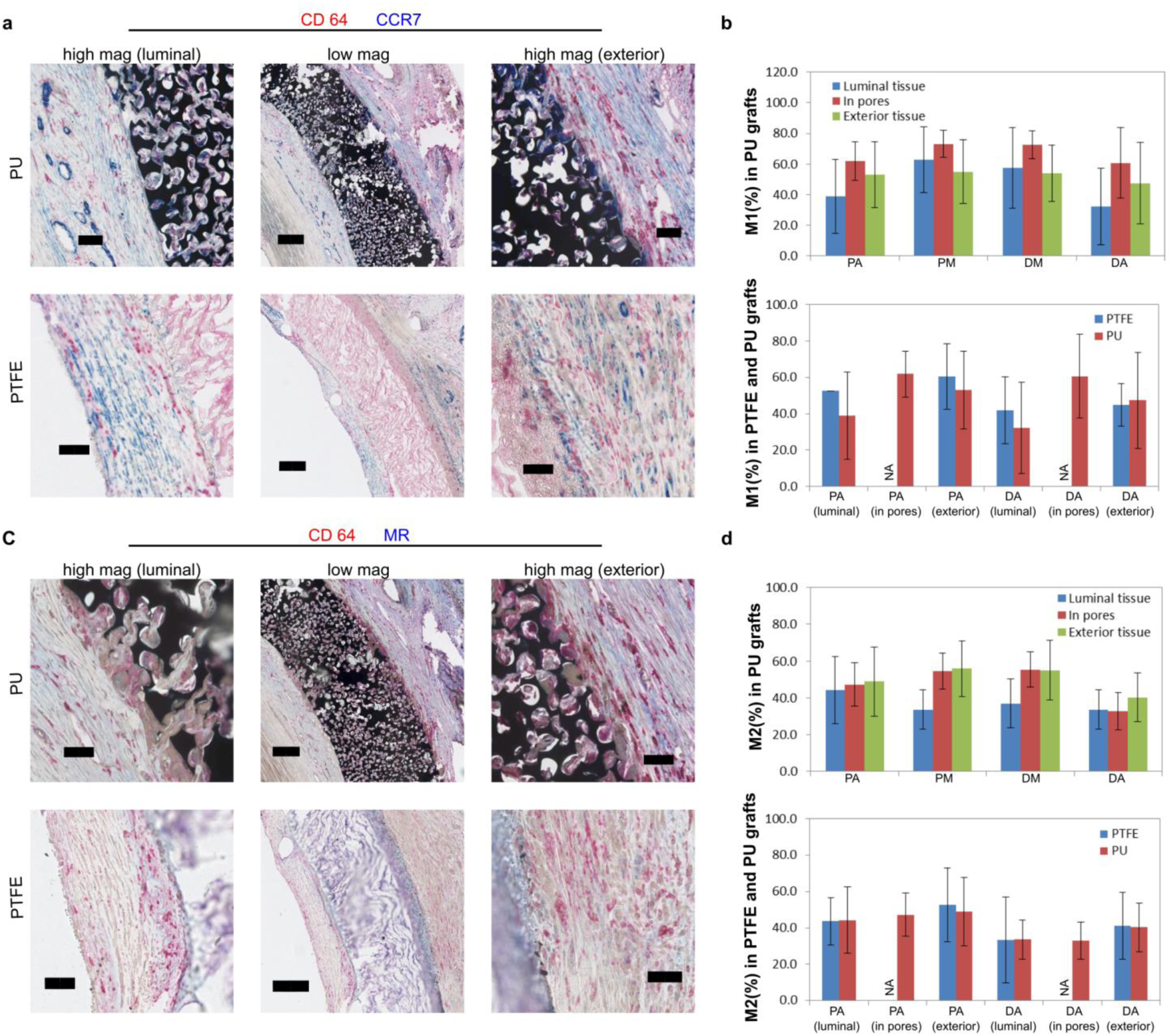
Immunohistochemical assessment of polarity of CD64+ cells. **a)** Representative micrographs of CD64 (a macrophage marker, in red) and CCR7 (an M1 marker, in blue) double stained cross-section of PU (top, PU material stained in black) and PTFE (bottom) grafts, in which low magnification micrographs (scale bars: 200 µm) are shown in the central column and high magnification micrographs (scale bars: 50 µm) of the luminal side (left column) and the exterior side (right column) are shown on the sides; **b)** M1 (%) polarity in different sections of PU grafts (top; PA: proximal anastomosis, PM: proximal mid-graft, DM: distal mid-graft, DA: distal anastomosis) and comparison of M1 (%) polarity between PTFE and PU grafts (bottom); **c)** Representative micrographs of CD64 (in red) and mannose receptor (MR, an M2 marker, in blue) double stained cross-section of PU and PTFE grafts (panel arrangement and scale bars are the same as in **a)**); **d)** M2 (%) polarity in different sections of PU grafts (top, acronyms are the same as in **b)**) and comparison of M2 (%) polarity between PTFE and PU grafts (bottom).

Quantifications of different regions (luminal tissue, tissue in pores, exterior tissue) of histological sections from different segments (PA, PM, DM, DA, see **Fig. 2b**, right) of PU grafts show a consistent trend of elevated level of M1 phenotype in pores compared to surrounding tissue in every segment (**Fig. 5b**, top). This is consistent with previous observations in small animal models.^10,11^ Given that M1 macrophages are able to secrete the highest level of proangiogenic factors compared to other phynotypes^26^, numerous M1 macrophages occupying the space inside the PU graft wall likely contribute to the improved angiogenesis in pores and endothelialization on the lumen. No consistent trend or difference of %M1 was observed between healed tissue surrounding PU and PTFE grafts (**Fig. 5b**, bottom). To identify Type 2 macrophages (M2), histological sections were double-stained with CD 64 (red) and mannose receptor (MR, blue) and cells positive for both markers were identified as M2 (purple, **Fig. 5c**). Specific staining was observed in healed tissue (in and around PU graft walls, around PTFE graft walls) and can be quantified, while unhealed tissue (blood clots in the interstices of PTFE grafts) mainly shows non-specific staining, and thus cannot be meaningfully quantified (**Fig. 5c**). Quantifications of M2 phenotype in PU grafts show a consistent trend towards higher %M2 in the exterior tissue compared to the luminal tissue (**Fig. 5d**, top), also consistent with observations by Sussman, et al.^11^ This trend is more pronounced in the mid-graft segments (PM, DM) compared to the anastomoses (PA, DA). The %M2 of tissue in pores of PU grafts mostly falls in the range between the exterior and luminal tissues (**Fig. 5d**, top). It was observed that, as healing progresses, the M2 marker increases over time.^27,28^ In light of this, the %M2 distribution observed in PU graft indicates that the healing may happen first at the exterior side, continue through the graft wall, and reach the lumen last (transmural healing). This would account for %M2 being higher in the exterior tissue that the healing process has been ongoing for a longer time compared to the luminal tissue. No consistent difference in %M2 is observed in healed tissues around PTFE and PU grafts (**Fig. 5d**, bottom). Overall, we propose that the advantage of the PU graft is that the porous structure can attract large numbers of macrophages to reside throughout the graft wall. These macrophages show the highest %M1 (proangiogenic) compared to surrounding tissues. At the same time, these macrophages maintain a relatively high %M2 (prohealing) comparable to exterior healed tissues, especially in the mid-graft regions. The abundance of M1 and M2 macrophages within the porous structure of PU grafts likely contribute to the improved healing observed in PU grafts.

### The PU material modulates FBGCs towards Type 1/Type 2 dual polarity

FBGCs were investigated through staining serial histology sections (**Fig. 6a**) with H&E (middle column), CD 64 and CCR7 (Type 1 marker, left column), CD 64 and MR (Type 2 marker, right column). MNGCs (pointed at by arrows) closely associated with graft materials (also known as foreign body giant cells, or FBGCs) were first identified by a pathologist in the H&E stained histological sections (**Fig. 6a**, middle column). The same cells (pointed at by arrows) were then found, respectively, in Type1 (**Fig. 6a**, left column) and Type 2 marker (**Fig. 6a**, right column) stained histological sections to determine their marker expression levels. In the representative images (**Fig. 6a**, top), all three FBGCs on the edge of the PU material are intensely stained with CCR7 (left) and positive for MR and CD 64 (right). In contrast, FBGCs on the edge of the PTFE material (**Fig. 6a**, bottom), although mostly positive for CD 64, are either only lightly stained with, or negative for CCR7 (left), and none are positive for MR (right).

**Figure 6.**
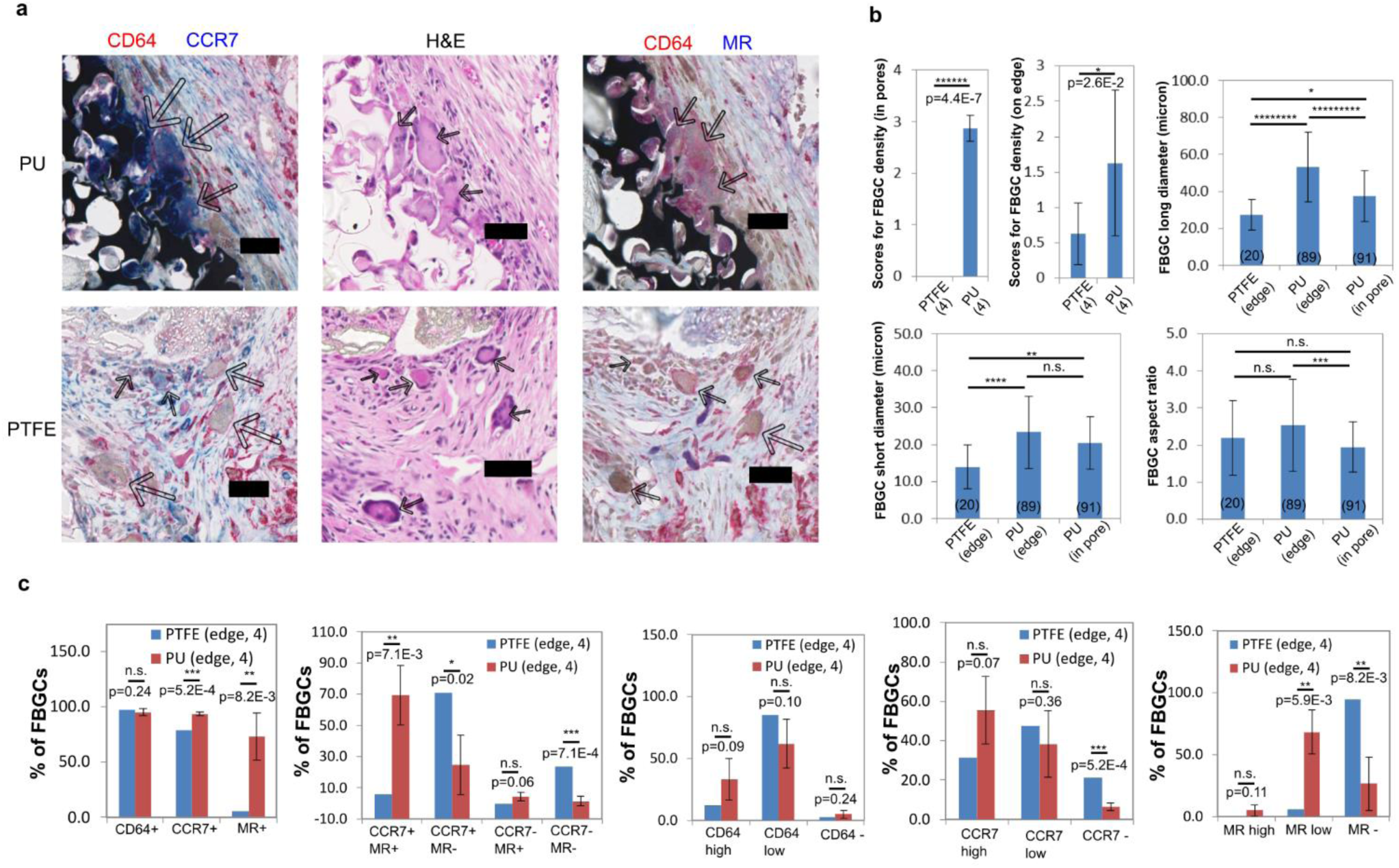
Assessment of foreign body giant cells (FBGCs) properties. **a)** Juxtaposition of same areas of three serial histological sections stained with CD64 (in red) and CCR7 (a Type 1 marker, in blue, left image), with H&E (middle image), and with CD64 (in red) and mannose receptor (MR, a Type 2 marker, in blue, right image), in which same FBGCs in each image are pointed out by arrows (scale bars: 50 µm); **b)** quantitative and semi-quantitative assessment (based on H&E staining) of FBGC density (top left two, numbers under graft labels show numbers of biological replicates (sheep)), size (top right and bottom left), and shape (aspect ratio, bottom right, numbers at the base of each bar show the number of FBGCs examined); **c)** quantitative assessment of immunohistochemical characteristics of FBGCs (in terms of percentage of FBGCs exhibiting certain characteristics). Numbers of sheep are shown in parentheses next graft labels. Percentages of FBGCs from PTFE grafts were calculated from all FBGCs found in all four sheep examined, due to the small number of FBGCs found in each sheep.

FBGCs identified by H&E were characterized semi-quantitatively or quantitatively (**Fig. 6b**). The density of FBGCs is significantly higher in the pores of PU grafts compared to PTFE grafts (p=4.4×10^-7^), with no FBGCs inside the latter due to pore size exclusion (**Fig. 6b**, top left). FBGC density on the edge of PU materials is also significantly higher than that of PTFE materials (pair-wise t-test, p=0.026, **Fig. 6b**, top middle). This indicates that the PU material is more inducive for FBGC formation, potentially due to the large numbers of macrophages it attracts. Next, the size (long axis, short axis) and shape (aspect ratio) of FBGCs on the edge of PU and PTFE grafts, and in the pores of PU material were examined (**Fig. 6b**, top right, bottom). In terms of long axis (**Fig. 6b**, top right), FBGCs on the edge of PU material are the largest (53.2±18.9 µm), FBGCs in pores of PU material are the second (37.5±13.9 µm), and those on the edge of PTFE material are the smallest (27.4±8.4 µm). The reduction in long axis from FBGCs on the edge of PU material (in contact with, or in close proximity of the material, but not subjected to the special confinement of pores) compared to those in pores is likely the result of spatial confinement. FBGCs on the edge of PTFE material, being the smallest, may reflect the material’s lack of attraction to macrophages, or immune inertness. The short axis of FBGCs shows a similar trend to long axis (**Fig.6b**, bottom left). In terms of aspect ratio (**Fig. 6b**, bottom right), FBGCs on the edges of PU and PTFE are comparable, while FBGCs in pores are significantly lower than FBGCs on the edge of PU. This suggests that FBGCs on edges can spread, while FBGCs in pores are spatially confined and not allowed to spread to the same extent. Lastly, the percentage of FGBCs expressing certain markers were counted (**Fig. 6c**). Close to 100% of FBGCs on edges of both materials are positive for CD 64 (**Fig. 6c**, left), indicating this is a relatively good marker for FBGCs. Higher percentages of FBGCs on the edge of PU are positive for CCR7 and MR, compared to PTFE (**Fig. 6c**, left, respectively, p(CCR7)=5.2×10^-4^ and p(MR)=8.2×10^-3^), indicating increases in both polarities of FBGCs associated with PU material.

FBGCs were further sorted into four groups based on their expression of (or lack of) markers: dual polarity giant cells (CCR7+, MR+, G1/2), type 1 giant cells (CCR7+, MR-, G1), type 2 giant cells (CCR7-, MR+, G2), and unpolarized giant cells (CCR7-, MR-, G0). The majority (69.4±19.2%) of FBGCs on the edge of PU are G1/2, which is significantly higher than those of PTFE (5.9%, **Fig. 6c**, 2^nd^ to the left, p=7.1×10^-3^). This suggests that the precision porous material induces a complex, or balanced, phenotype of both Type 1 and Type 2 characteristics in FBGCs. This is consistent with the observation in macrophages associated with such materials.^10^ This G1/2 phenotype may exert pro-healing effects. On the other hand, the majority (70.6%) of FBGCs on the edge of PTFE are G1, significantly higher than those of PU (24.8±19.1%, **Fig. 6c**, 2^nd^ to the left, p=0.02). This indicates that a simple Type 1 polarity may not lead to pro-regenerative healing. There are low percentages of G2 in FBGCs on the edge of both PTFE (0%) and PU (4.3±2.9%, **Fig. 6c**, 2^nd^ to the left). This suggests that a simple Type 2 response may not be common in the complex *in vivo* environment. The percentage of G0 on the edge of PTFE (23.5%) is higher than those of PU (1.5±3.0%, **Fig. 6c**, 2^nd^ to the left, p=7.1×10^-4^). This suggests the PTFE material is more immune-inert.

FBGCs were also counted according to their individual marker’s expression levels (high, low, negative, **Fig. 6c**, last 3 figures). The majority of FBGCs express low but distinguishable level of CD64 (**Fig. 6c**, middle). A lower percentage of FBGCs express high level of CD64 and an exceptionally small portion of FBGCs express no CD 64 (**Fig. 6c**, middle). FBGCs’ expression of CD64 does not vary significantly between the two materials. This shows CD64 to be a stable marker for FBGCs. Substantially higher percentage of FBGCs on the edge of PU express high level of CCR7 and significantly lower percentage of those express no CCR7 (p=5.2×10^-4^), compared to FBGCs on the edge of PTFE (**Fig. 6c**, 2^nd^ to the right). A similar percentage of FBGCs express low level of CCR7 in both materials (**Fig. 6c**, 2^nd^ to the right, p=0.36). The majority (94.4%) of FBGCs on the edge of PTFE are negative for MR, while 68.2±17.9% of FBGCs on the edge of PU express low level of MR (**Fig. 6c**, right). Overall, FGBCs on the edge of PU simultaneously up-regulate Type 1 and Type 2 markers. Expressions of markers do not vary between FBGCs on the edge of PU and FBGCs in pores of PU (**Fig. 3S**). This suggests that, although the confinement of the porous structure alters the size and shape of the FBGCs (**Fig. 6b**), it does not affect their polarity *in vivo*.

## Discussion

Our results support the hypothesis that precision porous PU grafts improve healing and endothelialization by recruiting a high number of macrophages and FBGCs, and modulating them towards complex, pro-healing phenotypes. We found that PU grafts are almost completely healed with tissue, both inside graft walls and along the lumen, while healing on PTFE grafts is limited to the vicinity of anastomoses. Overall endothelialization is significantly improved in PU grafts. Endothelialization in the middle region of PU grafts was observed. The precision porous structure of PU graft walls was infiltrated with nucleated cells, including mononuclear cells (mainly macrophages), FBGCs, and endothelial cells. All these cells reside within PU graft walls at significantly higher density compared to PTFE grafts. The interstices of PTFE graft walls were mainly filled with red blood cells (clots). High percentages of macrophages in pores of PU grafts showed high M1 (CCR7+) and M2 (MR+) characteristics. Immunohistochemical examination of FBGCs revealed that there exists a wide variety of phenotypes, including G0, G1, G2, and even G1/2. The majority FBGCs surrounding PU grafts are G1/2 (CCR7+MR+), indicating their complex, potentially pro-healing functions.

Remarkably, we demonstrated almost complete luminal coverage by cells with significantly enhanced endothelialization at 4 weeks (a time point strategically chosen for the well-established and robust immune response^29,30^) in a slow healing sheep model. This rapidity of healing in a sVG is unprecedented to our knowledge, and we believe this excellent healing is driven by the precision controlled 40 µm porous structures allowing for transmural cellular infiltration. A bilayer sVG without precision controlled porous structure took 6-24 months to achieve the healing visually similar to our grafts.^31^ In another study, coating sVG with fibronectin and stem cell homing factor achieved 48% of endothelial cell coverage (comparable to our PU grafts) in 3 months^32^ A biodegradable sVG showed good endothelialization in 6 months.^33^ With more sophisticated biological engineering, an *in vitro* tissue engineered decellularized graft revealed good endothelialization in 8-24 weeks.^34^

A decellularized artery was functionalized with proangiogenic factor to achieve endothelialization in 14 weeks.^35^ A small intestinal submucosa graft functionalized with heparin and vascular endothelial growth factor (VEGF) showed good endothelialization in one month.^36^ Our PU grafts achieved comparable or better endothelialization in more rapidly, without using sophisticated biologics. Thus, our approach is readily translatable to the clinic.

This proof-of-concept study is relatively short-term. To examine whether the rapid healing of our PU grafts translates into long-term clinical advantages, such as reduction of stenosis, thrombosis, and infection, longer-term implantation (90 days, 180 days) studies will be necessary next steps.

We discovered that, like macrophages, FBGCs exist in a spectrum of phenotypes. FBGCs in and around our PU grafts are mainly G1/2. This is consistent with previous observations in macrophages in the precision porous material.^10^ Given that a coordinated Type 1 and Type 2 response is crucial in promoting angiogenesis,^26^ the close association of the G1/2 with the pro-angiogenic effect of the porous material indicates that G1/2 may have a role in promoting angiogenesis similar to macrophages. This contrasts with the classical view that FBGCs were terminally differentiated and do not have diverse functions except giving off free radicals.^19^ FBGCs having a Type 2 characteristic (MR+) is in agreement with more recent *in vitro* studies. Specifically, IL-4 and IL-13 (two potent cytokines known to promote M2 polarization) induce macrophages to fuse and become FBGCs *in vitro*.^37,38^ Inhibition of MR activity blocks IL-4 induced FBGC formation *in vitro*.^39^ IL-1β (an M1 promoting cytokine) inhibits the formation of FBGCs *in vitro*.^40^ Based on these observations, FBGCs are suspected to be G2, not G1. We observed that, *in vivo*, there are few G2 FBGCs regardless of the graft material. Depending on the graft material, the FBGCs population are either dominated by G1 (PTFE) or G1/2 (PU). This discrepancy highlights the critical differences between *in vitro* and *in vivo* systems. A pure population of Type 1 (traditionally regarded as “pro-inflammatory”) or Type 2 (traditionally regarded as “pro-healing”) cells may be a product of the highly defined *in vitro* system where one cytokine is given at a time. *In vivo*, cells receive countless chemical and mechanical signals simultaneously and, as a result, may show more diverse and mixed phenotypes. Our results indicate that G1/2 may represent a complex, or balanced pro-healing phenotype *in vivo*.

In summary, we conducted a proof-of-concept study on the IMPRESSIVE graft in sheep carotid arteries. Compared to standard PTFE grafts, IMPRESSIVE grafts feature a precision controlled porous structure optimized for healing and improved dynamic compliance. After side-by-side implantations, PTFE grafts showed healing limited to the vicinity of anastomotic regions, while IMPRESSIVE grafts were almost completely infiltrated and covered by cells and healed tissue. Overall endothelialization is significantly improved and endothelialization at the middle region of IMPRESSIVE grafts was observed. Within the porous walls of IMPRESSIVE grafts, there reside higher densities of mononuclear cells, FGBCs, and endothelial cells, compared to PTFE grafts. The latter were mainly filled with red blood cells (probably clots). High percentages of macrophages in pores of the IMPRESSIVE graft showed M1 and M2 characteristics, indicating a coordinated pro-angiogenic response. We discovered that FBGCs, like macrophages, also exist in a spectrum of phenotypes. The dually activated phenotype G1/2 dominates the population of FGBCs associated with the pro-healing response to IMPRESSIVE grafts. The discovery of diverse phenotypes of FBGCs offers new insights into biocompatibility. To translate IMPRESSIVE to the clinic, longer-term implantation studies will be necessary. The IMPRESSIVE approach has wide applications for implantable medical devices and tissue engineering, in addition to pro-healing vascular grafts.

## Methods

### Fabrication of precision-porous vascular grafts

The method for making the polymer precision porous graft wall was reported in detail previously.^12^ Here we focus on describing the methods incorporating a mesh reinforcement and a gelatin sealant to make the full graft. A sterilized and roughened polyester knitted surgical mesh (TDA Textile Development Association Inc., PETKM 2006, PETKM 2008) was wrapped around a glass rod (6 mm in diameter) twice. The mesh-wrapped glass rod was inserted into a glass tube (8 mm in inner diameter) and thoroughly dried in a heated oven. The following steps for porous structure graft wall fabrication are as previously described (the polyurethane polymer formula PU 4344 was used).^12^ The porous graft wall was impregnated with gelatin (Sigma, Cat#1890-100G) by submerging the entire graft in 5% gelatin (w/v) solution and placing the solution at 60° C under vacuum for 5 minutes, followed by nitrogen purge of the vacuum. The impregnation was repeated 3 times. The graft was taken out to the solution and placed vertically at 4° C overnight to allow gelation. The graft was inverted, and the impregnation and cooling processes were repeated. The gelatin was crosslinked using EDC/NHS chemistry^41^ for 16 hours with the following conditions satisfied: crosslinking solution/gelatin solution(v/v)=50, NHS/EDC (molar ratio)=0.2, EDC/(-COOH in gelatin)=2. After the crosslinking reaction was completed, the graft was rinsed with DI water (2 hour ×3), 70% ethanal (2 hours ×9), and finally DI water (2 hours× 9). The graft was lyophilized, double packaged, and sterilized with ethylene oxide for 3 hours (Nelson Laboratories, Salt Lake City, UT). For each graft, a small section was cut and packaged separately and sterilized under the same condition for endotoxin and cytotoxicity (ISO10993). All of these small sections passed both tests before their separately packaged counterparts were implanted in sheep.

### Scanning electron microscopy (SEM) examination of porous structures

To examine the microporous structure of PU (without gelatin) and standard PTFE grafts, samples of both grafts were cut either on the circumferential or on the longitudinal direction, and then mounted on a metal stage using carbon tape with the cross-section, luminal surface, or the exterior surface facing up. The samples were sputter coated with gold using an MCM-200 Ion Sputter Coater for 60 seconds. The coating process was repeated once with samples rotated 180°. A silicon wafer etched with groves 10 µm apart (Planotec Test Specimen, Prod#615 Series, Wafer#E9613-06) was then mounted on the same stage and at the some height as the samples to serve as the calibration (the standard was not sputter coated). An SEM microscope (Nanoimages, Inc., SNE-3200M, equipped with Nanoeye software) was used to image the samples. To quantify the pore size and interconnect size of PU grafts, cross-sections of 5 different grafts were examined. Four high magnification images were taken from each cross-section. Ten pores and ten interconnects from each image were measured using Image J software.

### Dynamic compliance measurement

The dynamic radial compliance was evaluated by measuring the change in diameter of the graft under cyclic internal loading and in accordance with ISO 7198. Each graft was placed in-line to a circulatory flow-loop where unsteady flow was driven by a pulsatile pump (Harvard Apparatus, 1400, Boston, MA USA). The flow-loop was comprised of a flow-control valve to set resistance and an adjustable dynamic head which are used to modify the shape of the flow waveform. The internal pressure at the graft was measured by a pressure transducer (Omega Engineering Inc., Norwalk, CT, USA) directly upstream of the graft.

Experiments were conducted at pressure ranges of 50-90, 80-120, and 110-150 mmHg as well as 50-150 mmHg. All experiments were conducted at a cyclic frequency of 60BPM. A more detailed account for the compliance measurement method can be found in the supplement information.

### Implantation in sheep

All sheep experiments were approved by the University of Washington Animal Care and Use Committee (IACUC) and carried out in accordance with the National Institute of Health guide for the care and use of laboratory animals. Six adult female sheep (12-24 months in age, “mixed Willamette Valley” type) were purchased from Agna Farms (Salem, Oregon). All sheep tested negative for Q Fever before being shipped to the University of Washington were quarantined in house for at least 2 weeks before the surgery. Food was withheld 12-16 hours prior to the surgery, without controlling the water intake. Immediately prior to surgery, the neck area was shorn and shaved. The shaved area was cleansed with three consecutive rounds of betadine wash followed by 70% ethanol wash. After sheep were anesthetized, the eyes were lubricated. All surgical procedures were conducted within a draped, sterile field. Animal blood pressure, body temperature, and reflexes were continuously monitored during the procedure. Normal body temperature was maintained using a circulating water blanket. At the time of surgery, sedation and general anesthesia were induced and monitored by the Department of Comparative Medicine Veterinarian Services (DCM VS) staff. All administration of anti-stress drugs, analgesics, and anesthetics were supervised by attending UW veterinarians. A board-certified vascular surgeon conducted the implantation surgeries. Prior to the first incision, an IV bolus dose of 100-150IU/kg of heparin was given. Additional doses of 100IU/kg of heparin were given at 2 hours intervals until the surgery was completed. A dose of the antibiotic oxytetracycline was given. Lidocaine/bupivacaine were injected subcutaneously around the incision site prior to the first incision as a local anesthetic. In each surgery, each animal underwent bilateral interposition implantation of one PTFE and one PU graft (to allow side-by-side comparison of the two), each placed between a common carotid artery using interrupted suture technique.

Right and left side placement of grafts were randomized. Specifically, an incision approximately 7cm in length was made on one side of the neck to expose the common carotid artery. The artery was dissected free from surrounding tissue and encircled with vessel loops. Then a contralateral neck incision was performed and the contralateral common carotid artery was exposed in a similar fashion. Before the surgery, synthetic grafts were taken out of sterile packages and soaked in heparinized saline for at least half an hour. After clamping the artery, an interposition placement of synthetic graft was performed using either PTFE or PU graft (6mm ID, approximately 3.5 cm in length). The order and placement of the implanted grafts (PTFE vs PU in the right vs left) alternated between animals. The segment of native blood vessel replaced by vascular graft was harvested as control tissue. Anastomosis were performed end-to-end fashion using interrupted 6-0 polypropylene sutures. After completing the proximal and distal anastomosis, the vessel clamps were released, and patency of the graft was confirmed using a hand-held Doppler device. After patency of the first graft implant and animal homeostasis was confirmed, the contralateral grafting (with a different type of graft) was performed in a similar fashion.

After confirming patency on both sides, the incisions were irrigated with sterile saline and photographed. After hemostasis was confirmed, a layered closure was performed. Subcutaneous layers were closed with absorbable suture in a continuous pattern. The external skin closure was done with non-absorbable suture in an interrupted pattern. Each complete surgery was expected to take 3-4 hours. A total of 12 grafts (6 PTFE and 6 PU grafts) were implanted in 6 sheep. All sheep were allowed to recover for 4 weeks, during which internalized normal ratios (INR) between the undrugged prothrombin time and the drugged prothrombin time were monitored using the Roche CoaguChek XS system and controlled within a desired range of 1:2∼1:3.5 by daily oral administration of antiplatelet drug clopidogrel dosed in the range of 5.4-6.3mg/kg of body weight plus anticoagulant drug warfarin dosed in the range of 1.3-3.0mg/kg of body weight. Sheep were prepared for the harvesting surgery similar to the implantation surgery. After anesthesia was induced, ultrasound imaging was performed to confirm patency. Vascular grafts, together with a thin layer of surrounding tissue, the anastomoses and the connecting native blood vessel, were dissected free from the surrounding tissue. Photographs of the implants were taken. The sheep were euthanized by intravenous injection of a commercial euthanasia solution containing sodium pentobarbitol and sodium phenytoin. After death was confirmed, a suture loop was placed and tightened several centimeters above the distal anastomosis. Blood inside the grafts was flushed out by gentle injection of saline solution right beneath the suture loop. The native carotid arteries, the anastomosis, and grafts were harvested in block by transecting the carotid arteries several centimeters proximal and distal to the anastomosis. The grafts were immediately opened by cutting through the longitudinal direction using a pair of Metzenbaum scissors. The specimens were rinsed with heparinized saline so that only blood clots were retained on graft materials. The two types of grafts (PU and PTFE) from the same sheep were pinned down flat in saline and the luminal surfaces were photographed side-by-side. The pins were removed and the grafts were immediately fixed with freshly made 4% paraformaldehyde PBS solution for 48-72 hours. The grafts were processed, trimmed (as shown in **Figure 2b**) and embedded in paraffin.

### Basic histology and quantification

The whole grafts were cut in halves in a longitudinal direction and were routinely processed, paraffin embedded, and sectioned longitudinally at 5 µm thickness. 2-3 sections at least 200 µm (or 40 sections) apart from each other from each graft were examined by each type of staining. For H&E and Masson’s Trichrome, the staining and quantification protocols were described in detail previously.^12^ For calcification staining, 5 g of Alizarin Red S (abcam, ab146374) was added to 250 ml of deionized water.

Ammonium hydroxide solution was pipetted dropwise to the mixture under rigorous stirring to adjust the pH to 4.1-4.3. The solution was sterile filtered. Longitudinally sectioned slides were deparaffinized and rehydrated.

These slides were stained with freshly made Alizarin Red S solution for 2 minutes and excess dye was blotted. The histological sections were then dehydrated through acetone (20 dips), and acetone-xylene (1:1 solution, 20 dips), followed by clearing in xylene and mounting in a synthetic medium. Slides were scanned in brightfield with a 20x Plan Apo objective using the NanoZoomer Digital Pathology whole slides scanning system (HT-9600, Hamamatsu City, Japan). The slides were scored by a pathologist blinded to the hypothesis of the study (based on the scoring system shown in **Table S2**). A pairwise t-test was used to determine the statistical significance of the calcification scores (an infected sample and a sample with a suturing artifact were excluded).

### Immunohistochemistry and quantification

For endothelialization assessment, longitudinally sectioned whole graft samples were baked, deparaffinized, and rehydrated according to a protocol previously described.^12^ To stain the PU material black, an additional step, soaking the slides in 0.1% Sudan Black B solution (in 70% ethanol) was inserted in the rehydration process between the 95% ethanol wash and the 70% ethanol wash. Antigen retrieval was done by soaking the slides in boiling Tris-EDTA buffer (pH9.0) for 10 minutes, followed by cooling at room temperature for 20 minutes and rinsing in Tris buffered saline (TBS) for 5 minutes. To quench endogenous peroxidase activity, slides were submerged in 3% H2O2 TBS solution for 30 minutes, followed by 3 TBS rinses each for 5 minutes. Slides were then taken out of TBS and hydrophobic circles were drawn using a PAP pen around histological sections. Endogenous peroxidase and alkaline phosphatase (AP) activities were further quenched by incubating histological sections in BLOXALL solution for 20 minutes. After BLOXALL solution was removed, slides were rinsed in 3 rinses of TBS, each for 5 minutes. Nonspecific protein adsorption was blocked by incubating histological sections in 10% normal horse serum TBS solution (with 0.05% Tween-20) for 30 minutes. Slides were then incubated with TBS (with 2.5% normal horse serum and 0.05% Tween-20) diluent of anti-CD31 antibody (abcam, Cat#ab28364, from rabbit, initial concentration 0.013mg/ml, 1:300 dilution) and anti-Smoothelin antibody (R4A, abcam, Cat#ab8969, from mouse, initial concentration 1mg/ml, 1: 1000 dilution) overnight in a humidified chamber at 4°C. After antibodies diluent was blotted off, glass slides were rinsed in TBS solution for 5 minutes. Histological sections were incubated in ImmPRESS Duet Reagent (VECTOR, Ref#MP-7724) for 10 minutes, followed by two rinses in TBS each for 5 minutes. Positive staining of CD31 was visualized by incubating in Vector Blue substrate (VECTOR, Ref#SK-5300) until proper color was developed. Slides were then washed twice in TBS each time for 5 minutes. ImmPACT DAB EqV Substrate was applied to histological sections until proper color was developed. Slides were washed twice with TBS and coverslipped using VectaMount® AQ aqueous mounting medium (VECTOR, H-5501). The slides were stored flat in dark for 2 days and sealed using fingernail hardener before bubbles were formed. This procedure was applied to native carotid artery as a positive control and applied without primary antibody and with antibodies replaced by equal concentration isotypes as negative controls. In carotid artery controls, positive stains of CD 31 and smoothelin are clear and specific, almost no non-specific staining was observed in negative controls. However, this only remains true for CD 31 staining in grafts samples, where endogenous peroxidase (that cannot be quenched) from blood clots and newly healed tissue generate substantial background noise that obscures the smoothelin staining. Thus, only CD 31 staining was quantified. To quantify the CD 31 staining, slides were scanned in brightfield with a 20x Plan Apo objective using the NanoZoomer Digital Pathology whole slide scanning system (HT-9600, Hamamatsu City, Japan). Visiopharm software (Version 22.03.0.11565) was used to measure the full lumen length of each graft, continuous length of CD 31 positive lumen from the proximal end and the distal end, and length of CD 31 positive lumen apparently not connected to the proximal and distal end (mid-graft). The percentages of total endothelialized lumen in the total graft lumen length were calculated. Endothelial cell density in porous materials was assessed according to the method described previously.^12^

To assess macrophage and FBGC phenotypes, three serial histological sections in the circumferential direction were collected on three separate slides, where the middle sections were stained using H&E and the two sections on the sides were respectively double-stained immunohistochemically with CCR7 (Y59, abcam,Cat#ab32527, from rabbits, initial concentration: 0.347 mg/ml, 1:5000 dilution) and CD 64 (OTI3D3, abcam, Cat#ab140779, from mice, initial concentration: 1 mg/ml, 1:1000 dilution), or with mannose receptor (MR, abcam, Cat#ab64693, from rabbits, initial concentration: 1 mg/ml, 1:10000 dilution) and CD 64. To circumvent the problem of high endogenous peroxidase, two AP conjugated secondary antibodies, from AP Anti-Rabbit IgG Reagent Kit (Vector, MP-5401) and AP Horse Anti-Mouse IgG Polymer Kit (Vector, MP-5402) were sequentially applied after their corresponding primary antibody for color developments. A heat mediated antigen retrieval step was inserted in between the two color development steps to inactivate the first AP conjugated secondary antibody and activate the antigen (CD64) for the second primary antibody. This procedure takes advantage of the fact that CD64 requires antigen retrieval, while CCR7 and MR do not. Specifically, histological sections on the radial direction from PA, PM, DM, DA regions (as shown in **Figure 2b**) of PU grafts and PA, DA regions of PTFE grafts (sections from the PM and DM region would not remain attached to slides through the staining procedure due to lack of tissue integration and weak adhesion of the material to glass slides) were pretreated as described in the CD31 staining procedure (with the antigen retrieval step omitted). Appropriate diluent of the phenotype specific antibody (CCR7 for M1, or MR for M2) was applied to sections and incubated overnight. AP Anti-Rabbit IgG solution was applied to sections for 10 minutes after unbound primary antibody had been rinsed off. Proper color was developed using Vector Blue substrate. After rinsing, antigen retrieval was performed by placing slides in boiling citrate buffer (pH 6.0) for 10 minutes, followed by cooling for 20 minutes at room temperature.

BLOXALL treatment and nonspecific protein blocking were repeated. Proper diluent of macrophage-restricting antibody, CD 64, was applied to sections overnight, followed by TBS rinsing. AP horse anti-mouse IgG solution was applied to sections for 10 minutes, followed by rinsing with TBS twice each for 5 minutes. ImmPACT Vector Red from the Duet kit (VECTOR, Ref#MP-7724) was applied to sections until proper color developed. All slides were rinsed, coverslipped, sealed, and scanned as described above. This procedure was applied to both lymph node sections and PU graft sections as positive controls, and applied to these sections without primary antibodies, and with primary antibodies replaced by concentration-matched isotypes as negative controls. The clear and specific staining in positive controls and the absence of nonspecific staining in negative controls verifies the antibodies on both normal tissue and newly healed tissue on grafts. To make sure the first AP antibody was completely inactivated and did not cross-react with the second substrate, two more negative controls were established by 1) applying the procedure with only the second antibody (CD 64) replaced by isotype, and 2) applying the procedure with the first substrate (Vector Blue) omitted and second antibody replaced by isotype. Only blue staining was observed in the first negative control and no staining was in the second negative control, verifying the first AP antibody was fully inactivated after the first color development. To quantify the macrophage phenotypic responses, regions of interest (ROIs) were drawn around luminal collagenous tissue, porous materials, and exterior collagenous tissue using Visiopharm software (Version 22.03.0.11565). Colocalized staining of CD64 and the phenotype marker results in a purple hue, which was defined within the HIS color model as Purple=5-5.6 radians, Red=5.6-0 radians, Blue=2-5 radians. Within each ROI, the total area that stained red (positive for CD64 only), or A(red), and the total area that stained purple (positive for both phenotype marker and CD64), or A(purple), were measured. The overall percentage of certain phenotypic response, M(n)%, was calculated following the formular:

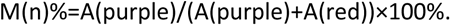

Where n=1 or 2 (the type of phenotypic response). To characterize MNGCs closely associated with both types of grafts (FBGCs), a professional pathologist, blinded to the hypothesis of the study, scored the density of FBGCs on the H&E stained histological section (the middle section of the three serial sections) based on a previously described method^12^, and annotated their locations. The FBGCs were treated as oval objects, where their long diameters and short diameters were measured, and aspect ratios were calculated. The same FBGCs annotated in H&E were located in the two double stained histological sections on the two sides and their marker expression levels were taken note of. The marker expression was depicted in three levels: high (intensely stained and/or widely distributed in the cell), low (faintly stained and distributed in small portion of the cell), and negative (no staining). The percentage of FBGCs expressing or co-expressing certain markers, as well as expressing certain level of markers were calculated. For PU grafts, at least 10-20 FBGCs from each graft were studied to extract the percentage information from individual sheep. An average and a standard deviation were calculated across different sheep. However, there were much fewer FBGCs found in PTFE grafts, thus we were not able to find enough cells in each graft to extract a percentage value. Instead, FBGCs examined from all the sheep were pooled together to calculate one percentage value. This overall value from PTFE grafts was set as the “theoretical” value in comparison with average value from PU grafts in one-sample t-tests to determine statistical significance.

### Statistics

Quantitative and semi-quantitative results are presented as mean ± standard deviation. Where it’s appropriate, statistical significance between groups were determined using unpaired two-sample student’s t tests for two groups (unless otherwise stated), and the ANOVA tests and the Bonferroni method for more than two groups. Statistical significance is defined by a p value (before Bonferroni modification) less than 0.05. Levels of statistical significance were denoted using the following system: n.s. means “not significant”; a single asterisk (*) denotes p<0.05; for more than one asterisks, the number of asterisks, n(*), indicates p<10^-n(*)^.

## Acknowledgement

The authors gratefully acknowledge a grant from the Northwest Kidney Centers to the University of Washington Center for Dialysis Innovation (CDI) for support during this study. We thank Paula and Steve Reynolds for their generous donation that supported the initiation of this study. We also thank KidneyX from the U.S. Department of Health and Human Services and the American Society for Nephrology, for a Phase II prize award to this project. Le Zhen would like to acknowledge A. Pat Miller Endowed Fellowship, and Graduate Assistant in Areas of National Need (GAANN) Fellowship for support of his PhD study on the project. Le Zhen would also like to express his gratitude to former undergraduate students Jason Dang, Melissa Gile, Isaac Lam, and Nicholas Zhen Hung Soo for their exploratory work under the GAANN Fellowship.

## Author Contributions

**L.Z.**: Conceptualization, data curation, formal analysis, Investigation, methodology, writing – original draft. **E.Q.**: Conceptualization, data curation, Investigation, methodology, writing – review & editing. **S.A.C.**: Investigation, methodology, project administration, writing – review & editing. **N.C.**: Investigation. **T.R.S.**: Investigation. **J.M.S.**: Methodology, investigation, writing – review & editing. **S.L.L.**: Methodology, investigation, software. **B.S.F.**: Methodology, investigation, software, writing – review & editing. **M.C.B.**: Methodology, investigation, writing – review & editing. **S.F.**: Investigation. **A.A.**: Methodology, supervision, writing – review & editing. **B.W.J.**: Methodology, supervision. **J.H.**: conceptualization, supervision, writing – review & editing. **B.D.R.** : conceptualization, methodology, supervision, writing – review & editing.

## Competing Interests

**B.D.R.**, **L.Z.**, and **J.H.** hold the patent (U.S. Patent No. 10,667,897) describing the IMPRESSIVE grafts studied in this paper.

