## Supplement Information for "Immunomodulatory Porous Regenerative Scaffolds for *in situ* Vascular Engineering"

**Optimization of the degradation rate for gelatin sealant.** The degradation rate of gelatin sealant was first optimized *in vitro*, followed by verification *in vivo*. Sterilely filtered 5% gelatin solution was pipetted into custom-made molds and refrigerated at 4 °C overnight to make 1-mm-thick gelatin sheets. Gelatin sheets were crosslinked (according to the procedure descripted in the method section) with n(EDC):n(-COOH)=1, 2, 4, making 1x, 2x, and 4x crosslinked gelatin. 6 disks punched out (using 6 mm biopsy punches) from each crosslinked density (made in three separated batches) were subjected to *in vitro* degradation test. Specifically, each disk was submerged in 1 ml of sterile Dulbecco's Modified Eagle Medium (DMEM) supplemented with 10% fetal bovine serum, 5% L-glutamine, 5% Penicillin-Streptomycin, and incubated at 36.5 °C with 5% $CO_2$. The culture medium was removed (daily for the first 7 days, and weekly afterwards until complete degradation) for scoring the level of degradation (based on **Table S1**) before replenishing with fresh medium. The degradation profile (**Fig. S1a**) shows that the time it takes to completely degrade *in vitro* is 7 days for 1x crosslinked gelatin, 30 days for 2x crosslinked gelatin, and 85 days for 4x gelatin. Given that *in vivo* degradation rate is likely to be faster, 1x crosslinked gelatin likely degrades too fast. 2x and 4x gelatin disks were implanted in mice for 21 days and stained with Masson's trichrome based on established protocols.[1] The stained histological sections show that 2x crosslinked gelatin completely degrades, is replaced by normal extracellular matrix and infiltrated by blood vessels within 21 days, while 4x gelatin only begins to degrade and is surrounded by classic foreign body capsule in the sample period of time (**Fig. S1b**). The level of degradation of 4x gelatin *in vivo* for 21 days is roughly equivalent to that of 40 days *in vitro*, from which it can be inferred that the *in vitro* degradation time is about twice of its *in vivo* counterpart. Based on these results, 2x crosslinked gelation was chosen as the sealant for the PU grafts. The pores of a PU (stained in black) graft can be efficiently sealed using 2x gelatin (stained in pink, **Fig. S1c**).

**Table S1.** The scoring system for the degree of degradation of gelatin disks.

| Score | Representative Image | Description |
|---|---|---|
| 0 | 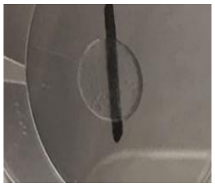 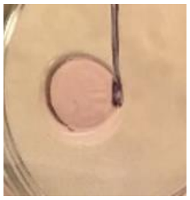 | Fully intact hydrogel with no swelling or degradation |
| 1 | 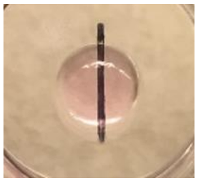 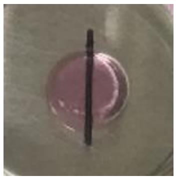 | Mildly swollen hydrogel (radius of disk looks larger than normal) |

| | | |
|---|---|---|
| 2 | 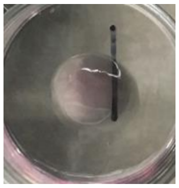 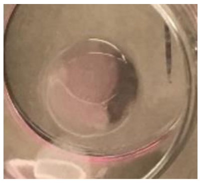 | Severely swollen hydrogel (disk grown substantially with water) |
| 3 | 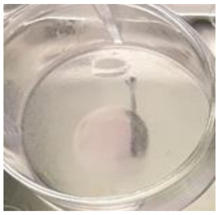 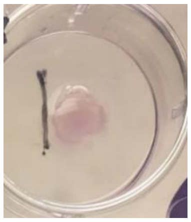 | Mildly degraded hydrogel (large portion of disk remaining) |
| 4 | 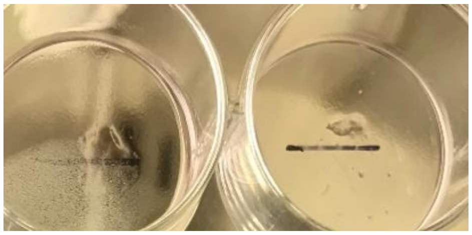 | Mostly degraded hydrogel (small portion of disk remaining) |
| 5 | 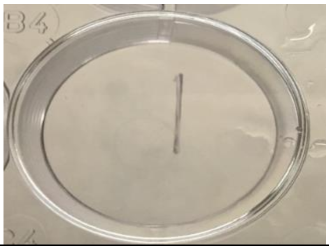 | Fully degraded hydrogel with nothing leftover |

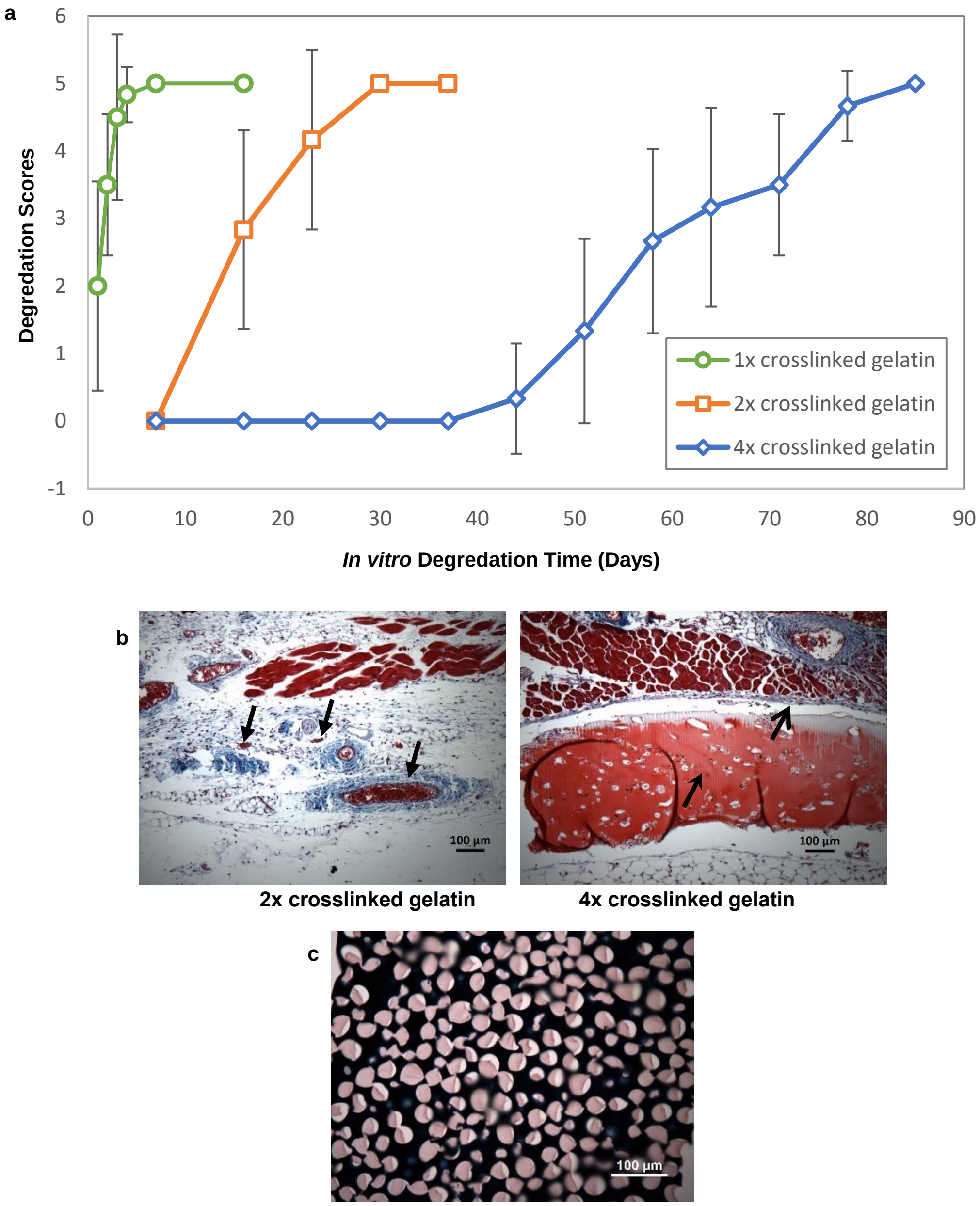


**Figure S1.** The optimization of the gelatin sealant. **(a)** The degradation profiles of gelatins with deferent crosslinking densities (1x, 2x, 4x) *in vitro*. **(b)** Masson's trichrome stained histological sections of 2x crosslinked gelatin (left) and 4x crosslinked gelatin (right) after 3 weeks subcutaneous implantation in mice. Arrows on the left point to blood vessels, open arrow on the right points to foreign body capsule, stealth arrow on the right points to undegraded gelatin. **(c)** A Masson's trichrome and Sudan Black B stained histological section of 2x gelatin (pink) sealed PU (black) graft.

**Dynamic compliance measurement (extended method).** The graft is connected to the flow-loop inside of an optically clear rectangular test-chamber that fixes the ends of the grafts in space such that the graft is free to only expand/contract radially. The distance between the graft connection points is held constant for each test. The diameter of the graft is optically measured using a high-speed camera (Phantom, V12, Vision Research, Wayne, NJ, USA). The camera is placed perpendicular to the front wall of the test chamber. An LED light panel is placed behind the optically clear test chamber, providing a high-contrast image of the graft. A diagram of the flow-loop and imaging configuration is shown in **Figure S2**. Before each test, a precision machined calibration rod with a diameter of 8mm is placed in the grafts position and a calibration image is acquired. The resolution of the imaging configuration is 12.7 microns/pixels. The camera capture (diameter measurement) and pressure measurement are synchronized, and measurements are acquired at 100 Hz. Data is collected for 10 seconds, and each experiment is repeated 3 times.

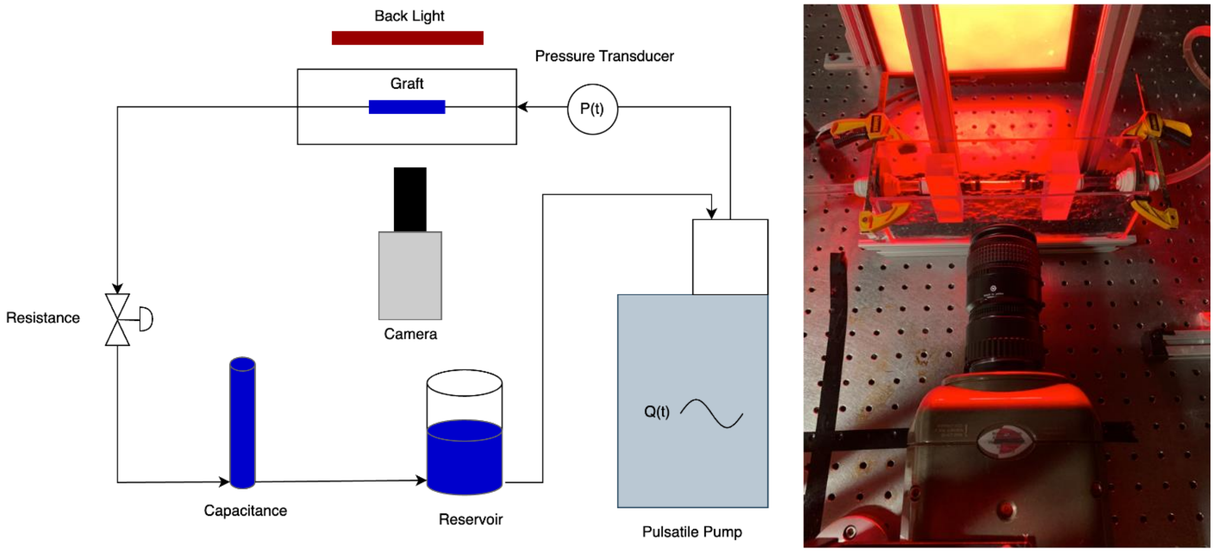


**Figure S2.** The setup for dynamic compliance measurement. (Left) A diagram of the flow-loop configuration. (Right) A photo showing the photo collection system.

To measure the diameter of the graft, a Sobel image filter is applied to each image in the sequence, providing a clear definition of the top and bottom edges of the graft. The diameter of the graft is measured as the distance between the two intensity spikes in the filtered image (graft edges). The diameter measurement is made at 10 stations across the length of the imaged graft. Dynamic compliance is then computed using the following equation:

$$C = \frac{\left[\left(\frac{D_{max} - D_{\min}}{D_{\min}}\right) x10^{4}\right]}{(P_{max} - P_{min})}$$

where $D_{min}, D_{max}, P_{min}, \& P_{max}$ represent the minimum and maximum diameters and pressures respectively. With the pressure and diameter measurements are synchronized, the compliance values are computed for each of the 10 flow cycles acquired and averaged in time. In **Figure 2a** (left), synchronized data and pressure measurements for a representative experiment are shown with the values of $D_{min}, D_{max}, P_{min}, \& P_{max}$ highlighted with red dots.

**Burst Testing**

To evaluate the structural integrity of the grafts, each graft is subjected to an internal pressure of 300 mmHg, which is 2.5 times of the normal systolic blood pressure of human. The graft is connected downstream of a compressed air line with a pressure regulator and variable opening control valve. A pressure transducer (Omega Engineering Inc., Norwalk, CT, USA) is placed upstream of the graft. The pressure regulator is set such that a maximum line pressure of 300 mmHg is achievable in the graft. The flow control valve is opened subjecting the graft to a uniform increase in pressure until 300 mmHg is achieved. The target pressure is held for 3 seconds and then the flow valve is closed and the pressure inside the graft allowed to return to ambient. The structural integrity (lack or presence of rupture) is evaluated via visual inspection of the graft and from the recorded pressure curve (a rupture would present as a sudden drop in pressure). The experiment is repeated 3 times for each graft sample.

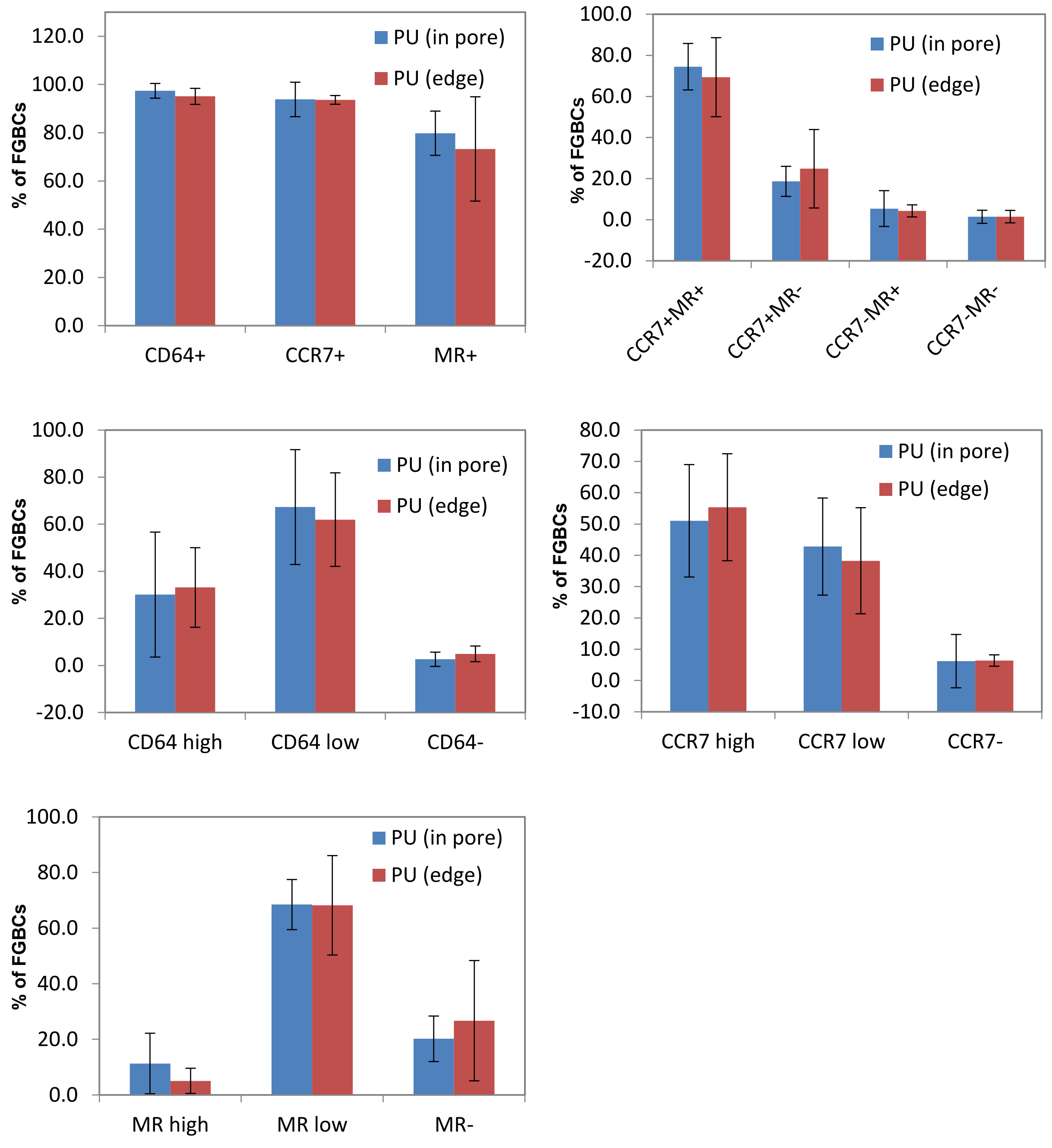


**Figure S3.** Percentages of FBGCs in pores of and on the edge of PU exhibiting certain immunohistochemical characteristics.

**Table S2.** Scoring system for semi-quantitative analysis of calcification in PU and PTFE grafts.

| Scores | Descriptions |
|---|---|
| 0 | absent |
| 1 | minimal, focal |
| 2 | minimal, multifocal, or mild focal |
| 3 | mild, multifocal, or moderate focal (regionally extensive) |
| 4 | moderate multifocal (multifocal to coalescing) or severe focal (effacing tissue architecture) |
